# Detection of live mycobacteria with a solvatochromic trehalose probe for point-of-care tuberculosis diagnosis

**DOI:** 10.1101/171553

**Authors:** M. Kamariza, P. Shieh, F. P. Rodriguez-Rivera, C. S. Ealand, B. Chu, N. Martinson, B. D. Kana, C. R. Bertozzi

## Abstract

Tuberculosis (TB) is the leading cause of death from an infectious bacterial disease. Poor diagnostic tools to detect active disease plague TB control programs and affect patient care. Accurate detection of live *Mycobacterium tuberculosis* (Mtb), the causative agent of TB, will improve TB diagnosis and patient treatment. We report that live mycobacteria can be specifically detected with a fluorogenic trehalose analog. We designed a 4-*N*,*N*-dimethylamino-1,8- naphthalimide-trehalose (DMN-Tre) conjugate that undergoes >700-fold fluorescence increase when transitioned from aqueous to hydrophobic environments. This enhancement occurs upon metabolic conversion of DMN-Tre to trehalose monomycolate and incorporation into the outer membrane. DMN-Tre labeling enabled the rapid, no-wash visualization of mycobacterial and corynebacterial species without nonspecific labeling of Gram-positive or –negative bacteria. DMN-Tre labeling was selective for live mycobacteria and was reduced by treatment with TB drugs. Lastly, DMN-Tre labeled Mtb in TB-positive sputum samples suggesting this operationally simple method may be deployable for TB diagnosis.

## Main Text

Tuberculosis (TB), caused by *Mycobacterium tuberculosis* (Mtb), is a serious global health challenge with an estimated 1.8 million deaths in 2015 (*1*). The increasing numbers of Mtb strains resistant to standard courses of treatment have exacerbated the global epidemic (*2*-*4*). The standard approach for rapid TB diagnosis, in high burden areas, is detection of Mtb in patient sputum using the color-based Ziehl-Neelsen (ZN) test, developed more than 100 years ago (*5*-*7*), or the fluorescent auramine-based Truant stain, first reported in 1938 (*8*). Both tests rely on the propensity of the mycobacterial outer membrane, also called the mycomembrane, to bind and retain hydrophobic dyes (*9*,*10*). The protocols require extensive processing to remove excess dye from debris and other bacteria while still retained by mycobacteria. Thus, the sensitivity of these tests varies widely (32% to 94%) depending on method and technician skill (*11*,*12*). Also, current diagnostic tests cannot distinguish live from dead mycobacteria (*13*) and therefore cannot report on treatment efficacy or guide clinical decision making in the face of rising drug resistance. New tools to advance sputum-based diagnostic accuracy, simplicity and specificity for live organisms are urgently needed.

Several groups, including ours, have observed that modified trehalose analogs, including fluorine- (*14*, *15*), azide- (*16*), alkyne- (*17*) or fluorophore-functionalized derivatives (*14*), can be metabolically incorporated into the mycobacterial outer membrane as trehalose mycolates. The process is enabled by the promiscuity of Mtb’s trehalose metabolic machinery, most prominently the antigen 85 complex (Ag85) that catalyzes mycolylation of trehalose (*18*) (Fig. 1A). Since trehalose mycolates are unique to the Corynebacterineae suborder, which includes pathogenic mycobacteria and corynebacteria but not canonical Gram-positive or negative organisms nor human hosts, the use of trehalose analogs for clinical detection of Mtb is an intriguing proposition. Indeed, fluorinated trehalose analogs have potential application for PET imaging of lung-resident Mtb (*19*,*20*). Likewise, we considered the possible use of fluorescent trehalose analogs for detection of viable Mtb as for point-of-care sputum smear tests, but the requirement for removing unmetabolized probe to eliminate background fluorescence proved to be a major impediment (*vide infra*). A trehalose probe whose fluorescence signal is specifically activated by metabolic incorporation into the mycomembrane would overcome such a limitation.

**Fig. 1.**
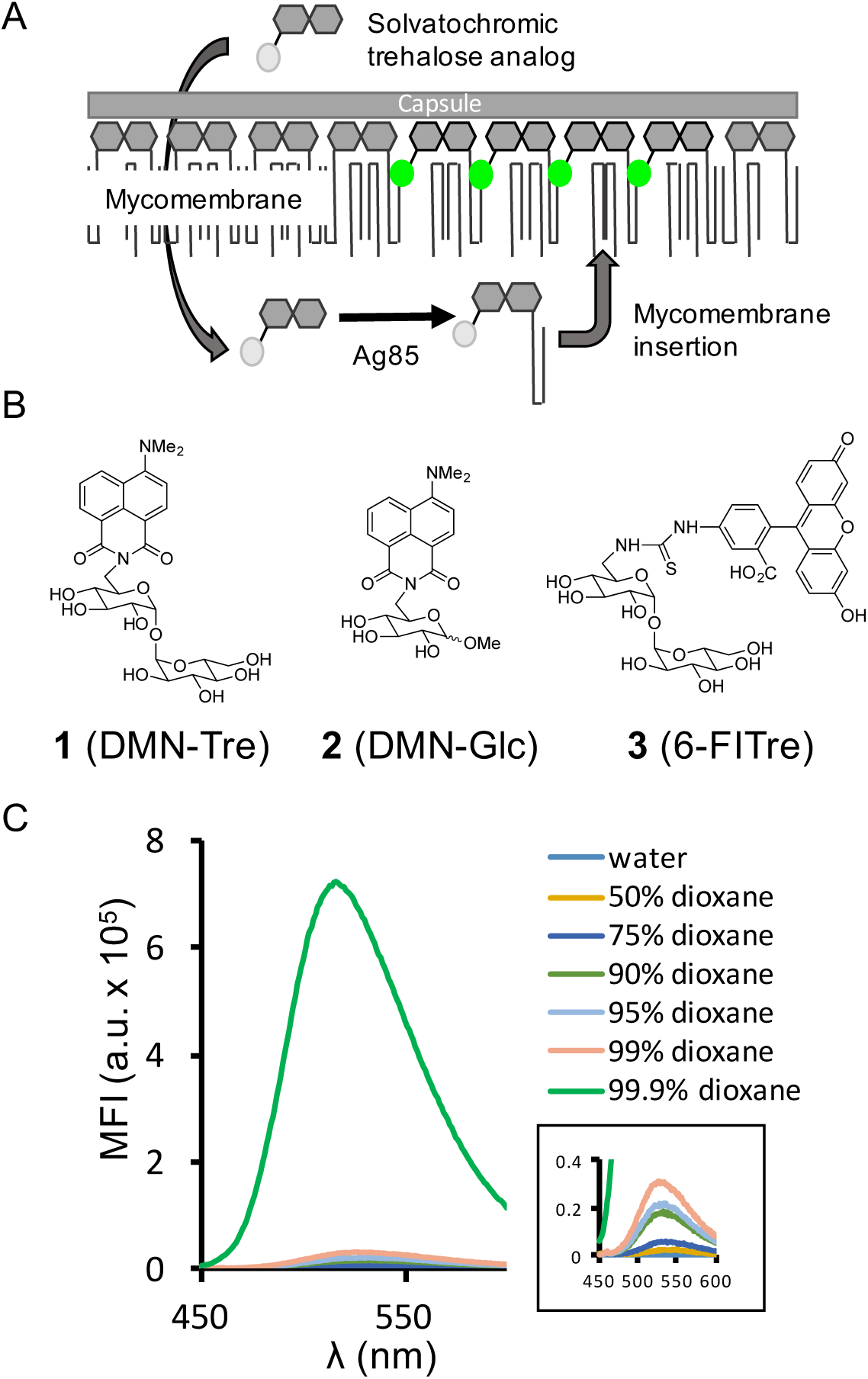
An environment-sensitive fluorogenic trehalose derivative for the detection of live mycobacteria. (A) Strategy for the detection of mycobacteria by metabolic incorporation of a trehalose analog functionalized with a solvatochromie probe into the mycomembrane. (B) Structures of trehalose-4-*N*,*N*-dimethylamino-l, 8-naphthalimide (DMN-Tre) and control compounds DMN-glucose (DMN-Glc) and 6-FlTre. (C) Fluorescence emission spectra of DMN-Tre in mixtures of dioxane:water. Inset shows enlargement of spectra in ≤ 99 % dioxane samples. MFI = Mean fluorescence intensity.

Here we show that trehalose conjugated to the solvatochromic dye 4-*N*,*N*-dimethylamino- 1,8-napthalimide (DMN), a reagent we call DMN-Tre, is metabolically incorporated into mycomembranes and enables fluorescent detection of mycobacteria in TB patient sputum samples. Unlike the classic ZN and auramine stains, DMN-Tre selectively labels live Mtb cells and this labeling is reduced by exposure to frontline TB drugs. This operationally simple method requires a single incubation step, with no washes, and may therefore powerfully complement current methods for TB diagnosis at the point of care.

We chose the environmentally-sensitive DMN dye based on work by Imperiali and coworkers’ observation that the molecule exhibits a dramatic fluorescence turn-on when transitioned from aqueous to organic solvents (*21*-*23*). Accordingly, we reasoned that metabolic mycolylation of a DMN-trehalose conjugate (DMN-Tre, **1**, Fig. 1B) and subsequent integration into the hydrophobic mycomembrane would activate fluorescence and enable detection of live Mtb cells without need for washing away unmetabolized probe (Fig. 1A). We synthesized DMN-Tre as well as two control compounds, DMN-glucose (DMN-Glc, **2**, Fig. 1B) and 6- fluorescein-trehalose (6-FlTre, **3**, Fig. 1B) (*24*), and confirmed that DMN-Tre’s fluorescence properties were similar with those reported for the free dye, including a ~700-fold enhancement in fluorescence intensity when dissolved in 99.9% dioxane versus water (Fig. 1C).

To evaluate DMN-Tre’s ability to label bacteria bearing mycomembranes, we tested several strains from the Corynebacterineae suborder. *Mycobacterium smegmatis* (Msmeg), *Mycobacterium marinum* and *Corynebacterium glutamicum*, each in exponential growth phase, were incubated with 100 μM DMN-Tre or the non-fluorogenic analog 6-FlTre for 2 hours then directly imaged without washing. Strikingly, we observed bright fluorescence labeling of all three species with DMN-Tre, with no discernible background fluorescence derived from free DMN-Tre in the surrounding solution (Fig. 2A). By contrast, cells labeled with 6-FlTre were obscured as expected by fluorescence of the probe in the surrounding solution. Extensive washing was required to remove nonspecifically-bound 6-FlTre from cells and their debris, rendering specifically labeled cells difficult to discern compared to those labeled with DMN-Tre. These images were acquired by using standard FITC/GFP filter sets, but an even brighter image can be obtained by excitation at 405 nm, closer to DMN’s excitation maximum (Fig. S3).

**Fig. 2.**
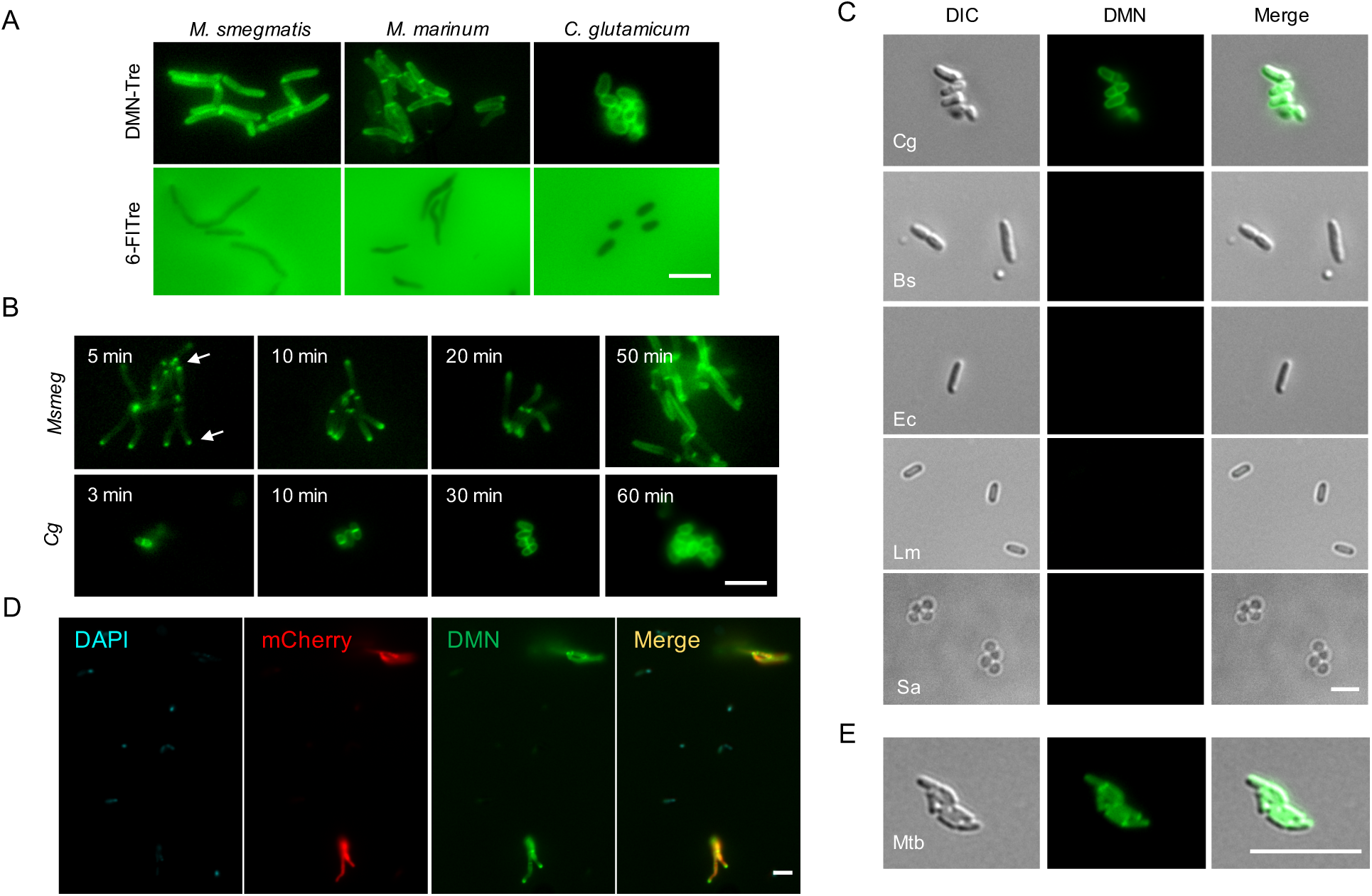
Specific, no-wash, detection of mycobacteria and corynebacteria using DMN-Tre. (A) No-wash imaging of *M. smegmatis* (Msmeg), *M. marinum*, and *C. glutamicum* (Cg) in the presence of 100 μM DMN-Tre or 6-FlTre for one doubling time. (B) Fluorescent labeling of Msmeg and Cg by DMN-Tre as a function of time. (C) Cg, *B. subtilis* (Bs), *E. coli* (Ec), *L. monocytogenes* (Lm), and *S. aureus* (Sa) cells were incubated with 100 μM DMN-Tre for 2 h then directly imaged. Background fluorescence with non-acid fast bacteria is minimal under no-wash imaging conditions. (D) Co-labeling of mCherry-expressing Msmeg, Bs, Ec, Lm and Sa under no-wash conditions. The mCherry Msmeg are preferentially labeled by DMN-Tre. (E) Mtb H37Rv cells were incubated with 100 μM DMN-Tre for 2 h followed by microscopy. Scale bar, 5 μm.

Next we performed a time course to determine the labeling kinetics. Msmeg and *C. glutamicum* (Cg) harvested during their exponential growth phase were incubated as above and imaged at various time points. As shown in Fig. 2B, labeling was already visible at the first time point – 5 min for Msmeg and 3 min for Cg. While Cg showed even cell surface labeling by the earliest time point analyzed, Msmeg showed fluorescence only at the polar regions, consistent with the polar growth model (*14*). This signal diffused across the cell length so that by 1 hour, Msmeg cells were uniformly labeled.

We performed a series of controls to confirm that DMN-Tre labeling results from metabolic conversion to trehalose mycolates within the mycomembrane rather than nonspecific insertion into the mycomembrane. DMN-Glc, which possesses the same dye but installed on glucose, did not label Msmeg, suggesting that the trehalose scaffold is key for the observed fluorescence signal from DMN-Tre treatment (Fig. S1A). As well, DMN-Tre labeling was reduced in the presence of excess trehalose, suggesting competition in the same biosynthetic pathway (Fig. S1B). We assessed DMN-Tre labeling of a panel of Msmeg trehalose transporter mutants (Fig. S1C) (*25*-*27*) and found that DMN-Tre incorporation was comparable to wild-type, suggesting that labeling occurs primarily through the Ag85 pathway, consistent with previous studies (*14*,*17*). Accordingly, the Ag85 inhibitor ebselen (*28*) decreased DMN-Tre labeling of Msmeg cells in a dose-dependent manner (Fig. S1D). Lastly, we analyzed purified glycolipids from DMN-Tre-labelled Cg by TLC (Fig. S2A) and mass spectrometry (Fig. S2B-D) and directly detected DMN-Tre-derived monomycolates.

DMN-Tre’s potential as a diagnostic tool for TB depends on its selectivity for mycobacteria among other bacterial species. We incubated canonical Gram-negative and – positive organisms (*Escherichia coli*, *Staphylococcus aureus*, *Listeria monocytogenes* and *Bacillus subtilis*) with DMN-Tre and, without any washing, observed no detectable labeling (Figs. 2C and S4). In the same experiment, Cg labeled brightly with DMN-Tre. This specificity is striking given the important role that free trehalose plays in these different bacterial species (*29*,*30*). Further, we combined mCherry-expressing Msmeg with these four bacterial species in a 1:10 mycobacteria/other bacteria ratio and incubated the mixture with both DAPI and DMN-Tre for one hour. We observed bright and specific labeling of only Msmeg (Fig. 2D). Lastly, we evaluated labeling of slow-growing pathogenic Mtb with DMN-Tre. Although labeling occurred more slowly compared to the faster growing Msmeg, we observed fluorescent cells within a 2- hour incubation period (Fig. 2E).

Current microscopy-based methods for TB diagnosis cannot distinguish live from dead mycobacteria. Given that DMN-Tre labeling depends on mycomembrane biosynthesis, we hypothesized that this method would be specific for live bacteria. Indeed, labeling was abrogated by heat killing Msmeg (Fig. 3A-B). We also evaluated the effect of TB drug treatment on labeling. Msmeg treated for 3 hours with a cocktail of ethambutol, rifampicin, isoniazid and SQ109, each at a dose at or above reported MICs, lost all detectable labeling with DMN-Tre (Fig. 3C-D). Similarly, drug-treated Mtb cells did not label with DMN-Tre (Fig. 3E, Fig. S5). In contrast, labeling with the commercial auramine-based mycobacterial staining dye (Fluorescent Stain Kit for Mycobacteria, Sigma-Aldrich cat. No. 05151) was unaffected by drug treatment (Fig. 3F).

**Fig. 3.**
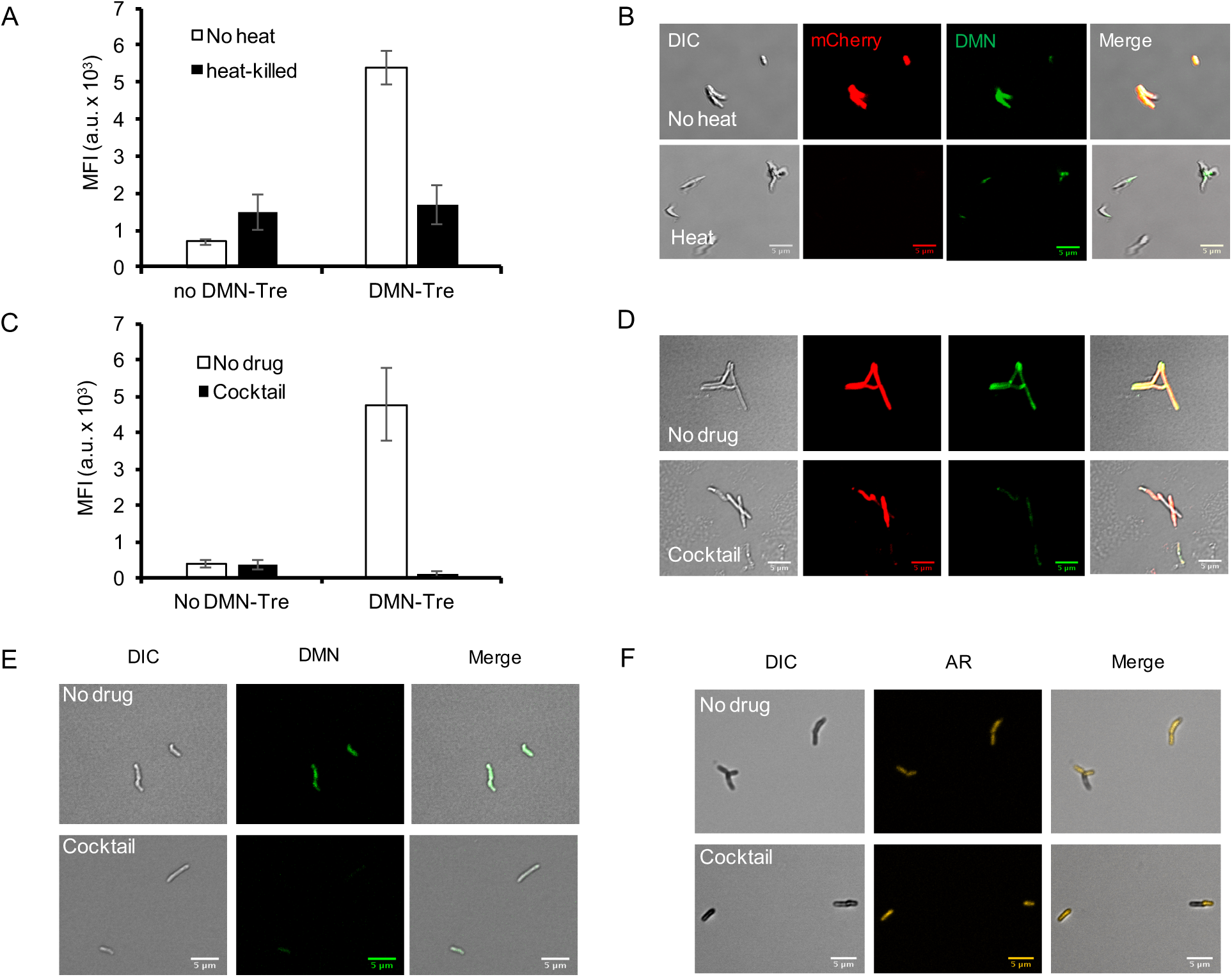
DMN-Tre labeling is selective for live mycobacteria and is diminished by TB drug cocktail. (A) Flow cytometry analysis and (B) no-wash imaging of control and heat-killed mCherry Msmeg cells after treatment with 100 μM DMN-Tre. Heat-killed cells were incubated in boiling water (temperature > 95 °C) for 30 minutes followed by DMN-Tre incubation. (C) Flow cytometry analysis and (D) no-wash imaging of control and drug-treated Msmeg cells in the presence of 100 μM DMN-Tre. Cocktail contents: 1 μg/mL ethambutol, 0.2 μg/mL rifampicin, 20 μg/mL SQ109 and 20 μg/mL isoniazid in 7H9 media. (E-F) Microscopy analysis of control and drug-treated Mtb cells labeled with (E) 100 μM DMN-Tre overnight or (F) stained with the auramine-based Fluorescent Stain Kit for Mycobacteria (Sigma-Aldrich, cat. no. 05151). Cocktail contents: 1 μg/mL ethambutol, 0.2 μg/mL rifampicin, 10 μg/mL SQ109 and 10 μg/mL isoniazid in 7H9 media. Scale bar, 5 μm.

To gain insight into the potential clinical utility of DMN-Tre, we sought to detect Mtb cells in TB-positive patient sputum samples. We obtained overnight sputum samples from 4 patients who were TB positive by either smear microscopy, GeneXpert or mycobacterial culture. The samples were decontaminated using a standard NaOH treatment, centrifuged to remove debris, and then incubated with DMN-Tre overnight (Fig. 4A). Fluorescent Mtb cells were readily apparent in all samples (Figs. 4B and S6).

**Fig. 4.**
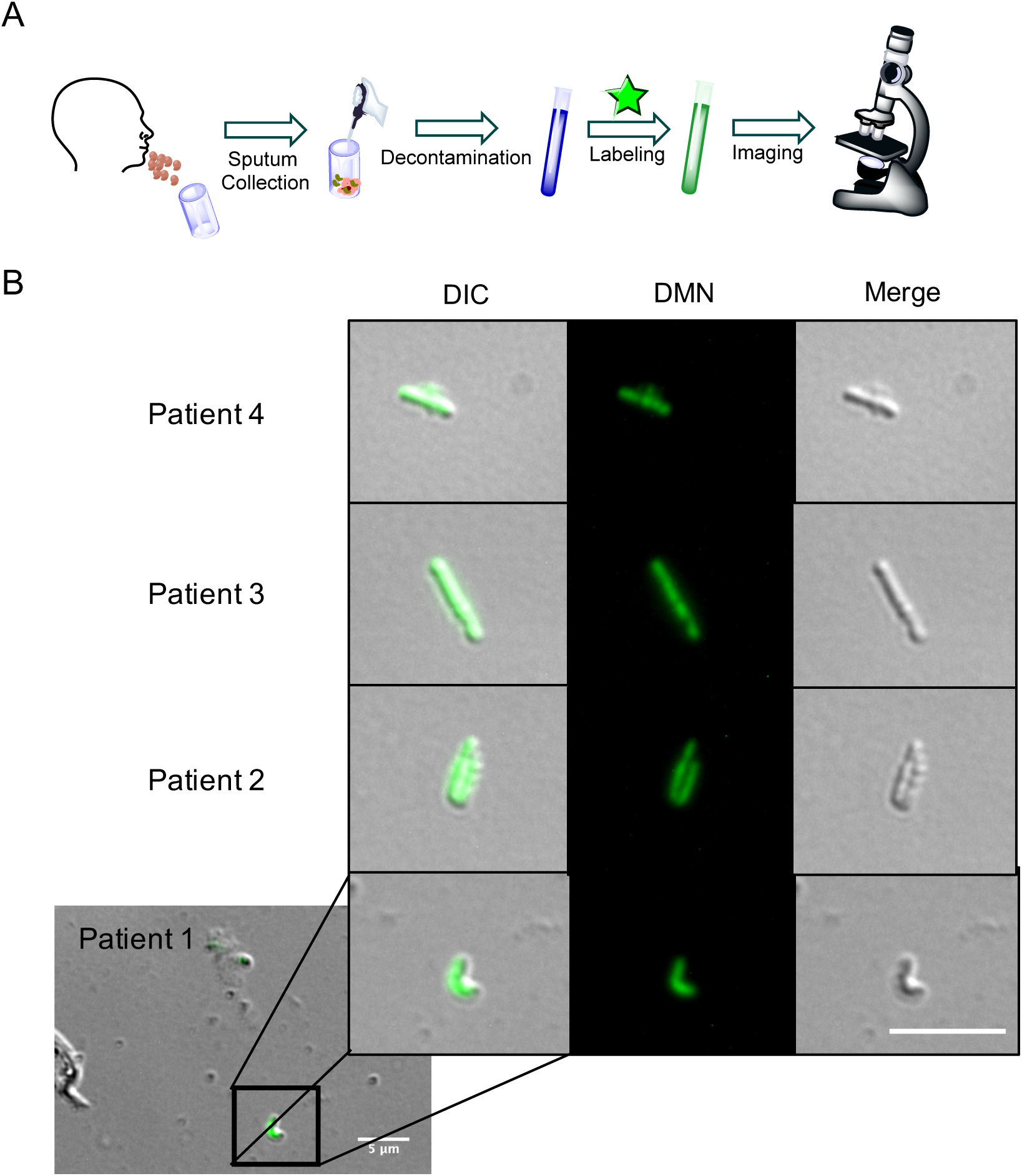
DMN-Tre detects Mtb in sputum sample from a TB-positive patient. (A) Illustration of sputum sample labeling protocol. (B) Samples were incubated with 1 mM DMN-Tre in 7H9 medium overnight. Inset shows enlargement of fluorescent cells. Panels depicts cells from 4 patient samples. Scale bar, 5 μm.

The World Health Organization has articulated the urgent need for new sputum tests for rapid TB diagnosis, particularly those that can report on drug susceptibility (*2*). Our data suggest that DMN-Tre labeling might fulfill this purpose. The method specifically targets a pathway in mycomembrane biosynthesis and therefore reports both on bacterial identity and vitality. DMN- Tre’s unique mode of fluorescence activation allows for an operationally simple procedure – a single incubation step. Notably, we found DMN-Tre to be very stable on the bench or in shipping containers for weeks at room temperature, and even in aqueous solution at 37 °C. Thus, DMN-Tre labeling holds much promise as reagent for TB research and diagnosis in low-resource settings.

## Acknowledgements

We thank F. Tomlin, D. Fox, Amanda McIvor and Nicole Narrandes for technical assistance, P. Robinson, C.J. Cambier and Melissa Chengalroyen for helpful discussions, and A. Iavarone (QB3/Chemistry Mass Spectrometry Facility, UC Berkeley) for assistance with mass spectrometry. Flow cytometry was performed in the shared FACS facility obtained with NIH S10 Shared Instrument Grant (S10RR027431-01). M.K. was supported by Stanford University’s Diversifying Academia, Recruiting Excellence fellowship. F. P. R.-R. was supported by a Ford Foundation Predoctoral Fellowship and UC Berkeley Chancellor’s Fellowship. B.D.K received an International Early Career Scientist Award from the Howard Hughes Medical Institute. Sputum collection and analysis was supported by the South African National Research Foundation to B.D.K. and C.S.E., the Centre for Aids Prevention Research in South Africa (CAPRISA) to C.S.E., and the Bill and Melinda Gates Foundation (Accelerator Grant: OPP1100182). This research was supported by the Bill and Melinda Gates Foundation (OPP115061) and NIH (AI051622) grants to C.R.B.

## Author contributions

P.S., M.K. and C.R.B. designed and led the study. M.K., P.S., F. P. R.-R., B.C. and C. S. E. performed laboratory experiments. C. S. E. and B. D. K. led the patient recruitment, patient sputum sample testing and analysis. M.K. and C.R.B. analyzed the data. M.K., P.S. and C.R.B. wrote and edited the manuscript which was approved by all authors.

